## Supplementary Table 2 for "Fibroblast-expressed LRRC15 suppresses SARS-CoV-2 infection and controls antiviral and antifibrotic transcriptional programs"

### Supplementary Table 2: Oligonucleotides for CRISPR activation sgRNA constructs

- All sgRNA sequences were taken from either the Human Genome-wide CRISPRa-v2 library (Addgene #83978) or Human CRISPR Activation Pooled Library (Calabrese P65-HSF) (Addgene #92379)
- Individual oligonucleotides were designed to be compatible with Esp3I- or BsmBI-v2-digested pXPR_502 (Addgene #96923).
  - To generate sense strand oligonucleotides, CACC was concatenated to the 5’ end of the sgRNA sequence taken from either of the two afore-mentioned libraries. If the sgRNA sequence did not start with a ‘G’, CACCG was concatenated tzo the 5’ end of the sgRNA sequence instead.
  - To generate antisense strand oligonucleotides, AAAC was concatenated to the 5’ end of the reverse complement of the sgRNA sequence taken from either of the two afore-mentioned libraries.
- All oligonucleotides were purchased from IDT

| **Oligonucleotide ID** | **Sequence (5’ → 3’)** | **Source** |
| --- | --- | --- |
| NTC sgRNA 1 - sense strand | CACCGCATCAGGAACCACGCGTCA | Human Genome-wide CRISPRa-v2 Library (Addgene #83978) |
| NTC sgRNA 1 - antisense strand | AAACTGACGCGTGGTTCCTGATGC | Human Genome-wide CRISPRa-v2 Library (Addgene #83978) |
| ACE2 sgRNA 1 - sense strand (Weissman) | CACCGGCGCCCAACCCAAGTTCAA | Human Genome-wide CRISPRa-v2 Library (Addgene #83978) |
| ACE2 sgRNA 1 - antisense strand (Weissman) | AAACTTGAACTTGGGTTGGGCGCC | Human Genome-wide CRISPRa-v2 Library (Addgene #83978) |
| ACE2 sgRNA 2 - sense strand (Weissman) | CACCGAGGAGAGGTAAGGTTCTCT | Human Genome-wide CRISPRa-v2 Library (Addgene #83978) |
| ACE2 sgRNA 2 - antisense strand (Weissman) | AAACAGAGAACCTTACCTCTCCTC | Human Genome-wide CRISPRa-v2 Library (Addgene #83978) |
| ACE2 sgRNA 3 - sense strand (Weissman) | CACCGGGCCATAAAGTGACAGGAG | Human Genome-wide CRISPRa-v2 Library (Addgene #83978) |
| ACE2 sgRNA 3 - antisense strand (Weissman) | AAACCTCCTGTCACTTTATGGCCC | Human Genome-wide CRISPRa-v2 Library (Addgene #83978) |
| ACE2 sgRNA 1 - sense strand (Calabrese Set A) | CACCGAGCCAATATAAAGTTCATCC | Human CRISPR Activation Pooled Library (Calabrese P65-HSF) (Addgene #92379) |
| ACE2 sgRNA 1 - antisense strand (Calabrese Set A) | AAACGGATGAACTTTATATTGGCTC | Human CRISPR Activation Pooled Library (Calabrese P65-HSF) (Addgene #92379) |
| ACE2 sgRNA 2 - sense strand (Calabrese Set A) | CACCGATATAAAGTTCATCCTGGAG | Human CRISPR Activation Pooled Library (Calabrese P65-HSF) (Addgene #92379) |
| ACE2 sgRNA 2 - antisense strand (Calabrese Set A) | AAACCTCCAGGATGAACTTTATATC | Human CRISPR Activation Pooled Library (Calabrese P65-HSF) (Addgene #92379) |
| ACE2 sgRNA 3 - sense strand (Calabrese Set A) | CACCGTTACATATCTGTCCTCTCC | Human CRISPR Activation Pooled Library (Calabrese P65-HSF) (Addgene #92379) |
| ACE2 sgRNA 3 - antisense strand (Calabrese Set A) | AAACGGAGAGGACAGATATGTAAC | Human CRISPR Activation Pooled Library (Calabrese P65-HSF) (Addgene #92379) |
| LRRC15 sgRNA 1 - sense strand | CAC CGA CAT GCA GGC ACT GCA CTG | Human CRISPR Activation Pooled Library (Calabrese P65-HSF) (Addgene #92379) |
| LRRC15 sgRNA 1 - antisense strand | AAA CCA GTG CAG TGC CTG CAT GTC | Human CRISPR Activation Pooled Library (Calabrese P65-HSF) (Addgene #92379) |
| LRRC15 sgRNA 2 - sense strand | CAC CGT ACC CTG TAG TGT CAG CCC | Human CRISPR Activation Pooled Library (Calabrese P65-HSF) (Addgene #92379) |
| LRRC15 sgRNA 2 - antisense strand | AAA CGG GCT GAC ACT ACA GGG TAC | Human CRISPR Activation Pooled Library (Calabrese P65-HSF) (Addgene #92379) |
| LRRC15 sgRNA 3 - sense strand | CAC CGT GTC CCG GGC TGA CAC TAC A | Human CRISPR Activation Pooled Library (Calabrese P65-HSF) (Addgene #92379) |
| LRRC15 sgRNA 3 - antisense strand | AAA CTG TAG TGT CAG CCC GGG ACA C | Human CRISPR Activation Pooled Library (Calabrese P65-HSF) (Addgene #92379) |
