## Supplementary Table 3 for "Fibroblast-expressed LRRC15 suppresses SARS-CoV-2 infection and controls antiviral and antifibrotic transcriptional programs"

### Supplementary Table 3: Primers for Next-Generation Sequencing of CRISPRa sgRNA from transduced cell gDNA

- Primer sequences adapted from:
  **Optimized libraries for CRISPR-Cas9 genetic screens with multiple modalities.** Sanson KR, Hanna RE, Hegde M, Donovan KF, Strand C, Sullender ME, Vaimberg EW, Goodale A, Root DE, Piccioni F, Doench JG. *Nat Commun. 2018 Dec 21;9(1):5416. doi: 10.1038/s41467-018-07901-8.* PubMed 30575746
- Primer regions:
  - P5/P7 flowcell attachment sequence
  - Illumina sequencing primer
  - Vector primer binding sequence
  - Stagger region / Barcode region

| **P5 Primers** | |
| --- | --- |
| **Stagger** | **Sequence** |
| **0** | AATGATACGGCGACCACCGAGATCTACACTCTTTCCCTACACGACGCTCTTCCGATCTCTTGTGGAAAGGACGAAACACC |
| **1** | AATGATACGGCGACCACCGAGATCTACACTCTTTCCCTACACGACGCTCTTCCGATCTGTCTTGTGGAAAGGACGAAACACC |
| **2** | AATGATACGGCGACCACCGAGATCTACACTCTTTCCCTACACGACGCTCTTCCGATCTAGTCTTGTGGAAAGGACGAAACACC |
| **3** | AATGATACGGCGACCACCGAGATCTACACTCTTTCCCTACACGACGCTCTTCCGATCTGCCTCTTGTGGAAAGGACGAAACACC |
| **4** | AATGATACGGCGACCACCGAGATCTACACTCTTTCCCTACACGACGCTCTTCCGATCTACAATCTTGTGGAAAGGACGAAACACC |
| **5** | AATGATACGGCGACCACCGAGATCTACACTCTTTCCCTACACGACGCTCTTCCGATCTTAGAGTCTTGTGGAAAGGACGAAACACC |
| **6** | AATGATACGGCGACCACCGAGATCTACACTCTTTCCCTACACGACGCTCTTCCGATCTCAGCAATCTTGTGGAAAGGACGAAACACC |
| **7** | AATGATACGGCGACCACCGAGATCTACACTCTTTCCCTACACGACGCTCTTCCGATCTTGAGACATCTTGTGGAAAGGACGAAACACC |

| **P7 Primers** | |
| --- | --- |
| **Index** | **Sequence** |
| 5 | CAAGCAGAAGACGGCATACGAGATCACGATGTGACTGGAGTTCAGACGTGTGCTCTTCCGATCTACCGACTCGGTGCCACTTTTTCAAG |
| 6 | CAAGCAGAAGACGGCATACGAGATCAGGCGGTGACTGGAGTTCAGACGTGTGCTCTTCCGATCTACCGACTCGGTGCCACTTTTTCAAG |
| 7 | CAAGCAGAAGACGGCATACGAGATTACAGCGTGACTGGAGTTCAGACGTGTGCTCTTCCGATCTACCGACTCGGTGCCACTTTTTCAAG |
