## Supplementary Table 4 for "Fibroblast-expressed LRRC15 suppresses SARS-CoV-2 infection and controls antiviral and antifibrotic transcriptional programs"

### Supplementary Table 4: Primers for RT-qPCR

| **Primer** | **Sequence (5’ → 3’)** | **Source** |
| --- | --- | --- |
| *GAPDH* Fwd | ACAACTTTGGTATCGTGGAAGG | Primerbank^1^ (Primerbank ID: 378404907c2) |
| *GAPDH* Rev | GCCATCACGCCACAGTTTC | Primerbank^1^ (Primerbank ID: 378404907c2) |
| *ACE2* Fwd | GGACCCAGGAAATGTTCAGA | Jia et al., 2020^2^ |
| *ACE2* Rev | GGCTGCAGAAAGTGACATGA | Jia et al., 2020^2^ |
| *LRRC15* Fwd | GCCTTTGGACAAGGCTATGC | Wang et al., 2018^3^ |
| *LRRC15* Rev | GAGCAGGTACACTCGCTAGG | Wang et al., 2018^3^ |
| *COL1A1* Fwd | CAAGACGAAGACATCCCACCA | This paper |
| *COL1A1* Rev | AGTAGCACCATCATTTCCACGA | This paper |
| *COL1A2* Fwd | GTGGCAGTGATGGAAGTGTG | Yu et al., 2018^4^ |
| *COL1A2* Rev | AGGACCAGCGTTACCAACAG | Yu et al., 2018^4^ |
| *COL6A1* Fwd | GACCTCGGACCTGTTGGGTAC | Yuri et al., 2020^5^ |
| *COL6A1* Rev | TACCCCATCTCCCCCTTCAC | Yuri et al., 2020^5^ |
| *COL6A2* Fwd | CAGGAGGTCATCTCGCCG | Piao et al., 2021^6^ |
| *COL6A2* Rev | GTTCTGCAGCTGGCTGATG | Piao et al., 2021^6^ |
| *COL6A3* Fwd | CCTAACCACATATGTTAGTGGAGGT | Dankel et al., 2020^7^ |
| *COL6A3* Rev | GAATGTCTCGCTTGCTCTCTG | Dankel et al., 2020^7^ |
| *COL12A1* Fwd | AAAGGGGAAAGGAAATCAGC | Mohassel et al., 2019^8^ |
| *COL12A1* Rev | TCACAGCATCTGTCTCTACTGGT | Mohassel et al., 2019^8^ |
| *IFIT1* Fwd | GGAATACACAACCTACTAGCC | Li et al., 2019^9^ |
| *IFIT1* Rev | CCAGGTCACCAGACTCCTCA | Li et al., 2019^9^ |
| *IFIT3* Fwd | TGAGGAAGGGTGGACACAACTGAA | Li et al., 2019^9^ |
| *IFIT3* Rev | AGGAGAATTCTGGGTTGTTGGGCT | Li et al., 2019^9^ |
| *MX1* Fwd | AGGACCATCGGAATCTTGAC | Ortiz et al., 2020^10^ |
| *MX1* Rev | TCAGGTGGAACACGAGGTTC | Ortiz et al., 2020^10^ |
| *OAS1* Fwd | GCGCCCCACCAAGCTCAAGA | Li et al., 2019^9^ |
| *OAS1* Rev | GCTCCCTCGCTCCCAAGCAT | Li et al., 2019^9^ |
| *OAS2* Fwd | ACCCGAACAGTTCCCCCTGGT | Li et al., 2019^9^ |
| *OAS2* Rev | ACAAGGGTACCATCGGAGTTGCC | Li et al., 2019^9^ |
